# Long-term Pannexin 1 ablation promotes structural and functional modifications in hippocampal neurons through the regulation of actin cytoskeleton and Rho GTPases activity

**DOI:** 10.1101/2021.11.03.467134

**Authors:** Carolina Flores-Muñoz, Francisca García-Rojas, Miguel A. Perez, Odra Santander, Elena Mery, Daniela Lopez-Espíndola, Arlek M. Gonzalez-Jamett, Marco Fuenzalida, Agustín D. Martinez, Álvaro O. Ardiles

## Abstract

Enhanced activity and overexpression of Pannexin 1 (PANX1) channels contribute to neuronal pathologies, such as epilepsy and Alzheimer’s disease (AD). In the hippocampus, the PANX1 channels ablation alters glutamatergic neurotransmission, synaptic plasticity, and memory flexibility. Nevertheless, PANX1-knockout (KO) mice still preserve the ability to learn, suggesting that compensatory mechanisms work to stabilize neuronal activity. Here, we show that the absence of PANX1 in the adult brain promotes a series of structural and functional modifications in KO hippocampal synapses, preserving spontaneous activity. Adult CA1 neurons of KO mice exhibit enhanced excitability, complex dendritic branching, spine maturation, and multiple synaptic contacts compared to the WT condition. These modifications seem to rely on the actin-cytoskeleton dynamics as an increase in actin polymerization and an imbalance between Rac1 and RhoA GTPase activity is observed in the absence of PANX1. Our findings highlight a novel interaction between PANX1, actin, and small Rho GTPases that appear to be relevant for synapse maintenance as a long-term compensatory mechanism for PANX1 deficiency.

## Introduction

Pannexin 1 (PANX1) is a heptameric channel that enables the movement of ions and small molecules between intracellular and extracellular compartments to contributing to paracrine communication in mammalian cells (*1*–*5*). PANX1 channels exhibit two modes of activity with different properties of ion and solute permeability(*6*). At negative potentials, PANX1 channels show a constitutive small pore ion channel activity characterized by low conductance, slightly anion selectivity (chloride ions) and outwardly rectifying currents (*7*–*10*), while at depolarizing potentials, these channels exhibit a large pore conformation with high conductance mediating a non-selective ionic flux responsible for the permeation of ATP and other metabolites. ATP released by PANX1 channels is triggered by intense or chronic neuronal activity (*11*, *12*), high concentrations of external potassium (*13*), mechanical stress (*14*), low oxygen conditions (*15*, *16*), ionotropic and metabotropic receptor signaling (NMDAR (*11*, *17*), P2X7R (*18*), α1AR (*19*)), and caspase-dependent cleavage of its carboxy-terminal (*20*, *21*). However, it remains unclear the precise contribution of PANX1 to neuronal function under resting activity.

Aberrant PANX1 activity has been implicated in several conditions affecting CNS (*4*, *22*), including ischemia (*23*, *24*), epilepsy (*25*–*27*), and Alzheimer’s disease (*28*). Nevertheless, PANX1 channels also mediates physiological processes in the central nervous system (CNS). In the adult mouse brain, PANX1 absence or blockade increases the excitatory synaptic transmission and modifies the induction of synaptic plasticity (*29*, *30*). During embryonic and early postnatal development, PANX1 ablation promotes neurite outgrowth and dendritic spine development, and the formation of network ensembles (*31*, *32*),. Thus, the fine regulation of Panx1 activity and expression could be a critical aspect for the proper functioning of neurons and brain circuits and is intriguing to know the long-term effect of PANX1 ablation on both processes underlying structural plasticity.

It is known that the polarized and highly branched morphology of neurons is crucial to establish synaptic contacts and hence neural circuits. In fact, plastic changes in neuronal circuits are believed to be the basis for high-order brain functions such as cognition (*33*), and perturbations in their regulation are associated with synaptopathies (*34*). Most of the excitatory synapses are localize on dendritic spines, highly dynamic structures exhibiting the capacity to change their morphology and density in response to neural activity(*35*). Thus, the synapses are constantly subjected to activity-dependent plasticity to store information. Consequently, they are simultaneously compensated to avoid instability in the circuits preserving their plasticity and cognitive abilities. In this regard, homeostatic regulatory changes occur at the cell and circuit level to adapt neural properties and their molecular components to provide a balanced control of synaptic strength (*36*). Several mechanisms have been reported, such as metaplasticity, synaptic scaling, and intrinsic plasticity (for review see (*37*–*40*)), which can be executed through a variety of cellular and molecular processes that differ depending on the development state and the initial mechanism of activity perturbation (*41*). Previously was reported that the lack of PANX1 activity in the adult hippocampus produces metaplastic changes in the induction threshold for LTP and LTD (*29*). However, this change was revealed under activity demand, and it is unknown if PANX1 ablation elicits additional modifications to preserve neuronal activity at resting conditions. Interestingly, a recent study reported the early postnatal PANX1 ablation prevent the homeostatic adjustment of presynaptic strength upon chronic inactivity.

Here, we show that PANX1 deletion in the adult mouse brain promotes several functional and structural modifications in hippocampal neurons without alteration in spontaneous activity. These modifications include enhanced excitability, a higher dendritic arborization, and spine maturation, compared to the wild-type condition. Furthermore, KO neurons also exhibited an increased size of the readily releasable pool (RRP) and the postsynaptic density (PSD) and an increased number of synaptic contacts at the ultrastructural level. However, the estimation of a multiplicity index revealed that the functional sites of release or functional contacts were lower in KO neurons. Since we observe an enhanced expression of actin-related proteins and the activated form of Rac1 along with augmented F-actin content in hippocampal tissue from KO mice, all these evidences strongly suggest that Panx1 plays a “stabilizing” role in neuronal function and morphology by modulating the actin dynamics in a mechanism involving the Rho-GTPase family

## Results

### PANX1 ablation increases neural excitability but preserves spontaneous release

We determine whether PANX1 deficiency elicits adaptations in synaptic strength under basal conditions by using whole-cell current- and voltage-clamp recordings in hippocampal neurons from KO and WT littermates (Fig. 1). First, we examined the effects of PANX1 ablation on the intrinsic electrical properties of hippocampal pyramidal neurons (Table 1). We observed that KO neurons displayed a lower threshold for action potential discharge (Fig. 1, A and B; WT versus KO *p=0.0430, Man-Whitney test). These changes in excitability seem to be not mediated by changes in passive neuronal properties as WT and KO neurons exhibited similar resting membrane potential, input resistance, and whole-cell capacitance (Table 1). Since passive properties were unchanged, we explored for modifications in voltage-gated ionic currents (Fig. 1, C and D). No apparent differences in I-V curves were observed (Fig. 1D), although we cannot rule out that selective ionic currents might be different between groups. Next, we used two approaches to ask whether the variations in excitability upon PANX1 ablation might affect the synaptic transmission. First, we looked at basal synaptic transmission mediated by AMPA and NMDA receptors by generating input-output curves by measuring extracellular field potentials at varying stimulus intensity (Fig. 1E to L). The slope and the fiber volley amplitudes of AMPA-and NMDA-mediated field excitatory postsynaptic potentials (AR-and NR-FP) were similar, suggesting no obvious changes in basal synaptic transmission between KO and WT mice. However, it is noteworthy that evoked AR- and NR-FP from KO neurons exhibited an increased tendency to show population spikes at growing intensities (Fig. 1, H and L), revealing latent greater excitability of the CA1 KO neurons. Next, to test any possible modification in basal synaptic activity, we record spontaneous and miniature excitatory postsynaptic currents (sEPSCs and mEPSCs). Accordingly, we observed that either the frequency or the amplitude of sEPSCs were similar between groups, indicating no apparent effect on basal glutamate release (Fig. 2, A to D). Astonishingly, we found that the absence of PANX1 alters the amplitude of mEPSCs (Fig. 2, E to H). PANX1-KO mice exhibit a higher amplitude of mEPSCs compared to WT (WT versus KO *p=0.0104, Man-Whitney test), but the frequency of mEPSCs was unaltered between groups. We then estimated a multiplicity index, an approximation to estimate the number of releasing sites and infer about synaptic contacts (*42*, *43*) (see methods). Interestingly, we found that the multiplicity index in KO neurons was significantly lower than that observed in WT (Fig. 2, G and H), indicating that KO neurons establish less functional contacts or possess fewer releasing sites.

**Fig. 1.**
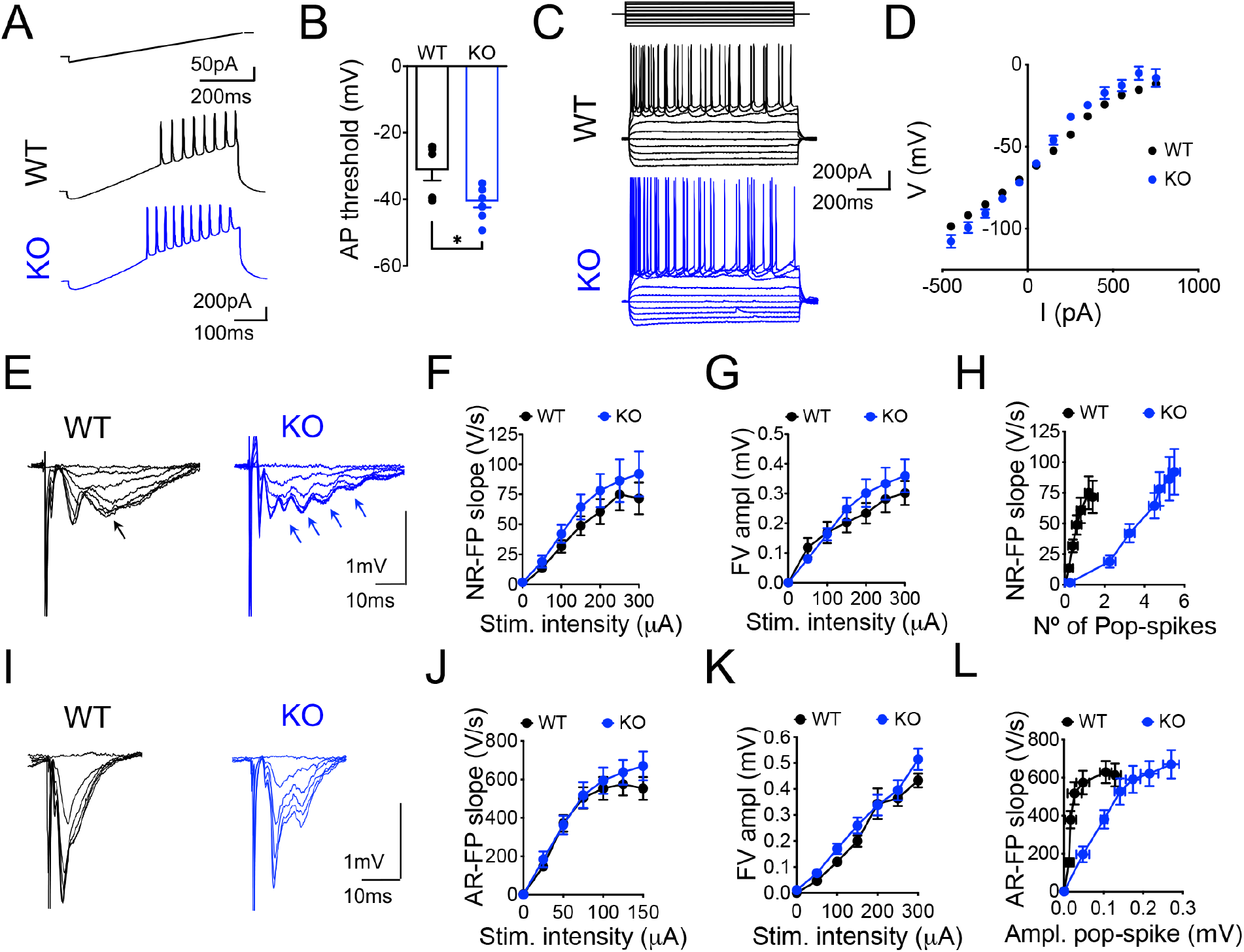
Enhanced excitability but normal glutamatergic synaptic transmission in hippocampal CA1 neurons from PANX1 KO mice. (**A**) Representative traces of membrane potential recordings in response to a current ramp. (**B**) Action potential threshold. (**C**) Representative traces of membrane potential changes in response to current steps. (**D**) Current-voltage curves (**E**) Representative traces of input-output curves of pharmacological isolated NMDAR fEPSP (NR-FP) and analysis of the slope (**F**), fiber volley (FV) amplitude (**G**) and plots of NR-FP slope versus the number of pop spikes (**H**). Representative traces (**I**) of input-output curves of AMPAR fEPSP (AR-FP) and analysis of the slope (**J**), FV amplitude (**K**), and plots of AR-FP slope versus the amplitude of the pop spike (**L**).

**Fig. 2.**
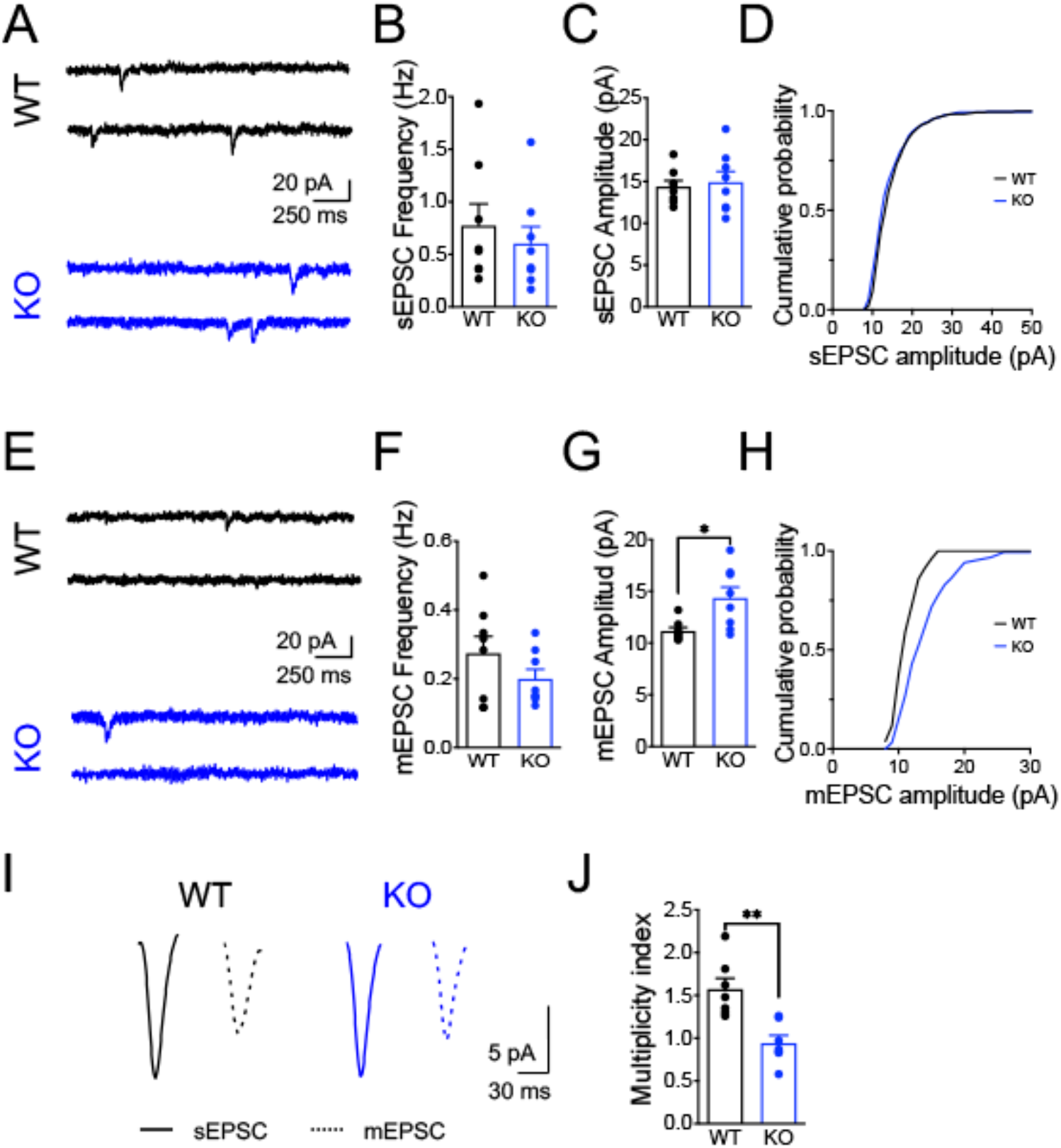
Increased mEPSC amplitude but normal spontaneous release and reduced number of releasing sites in PANX1 KO CA1 neurons. Representative traces (**A**) and analysis of sEPSC frequency (**B**) and amplitude (**C**) in CA1 neurons of wild type (WT, black) and PANX1-KO (KO, blue) mice. (**D**) Cumulative probability plots of the sEPSC amplitude distribution. Representative traces (**E**) and analysis of mEPSC frequency (**F**) and amplitude (**G**) in CA1 neurons of wild type (WT, black) and PANX1-KO (KO, blue) mice. (**H**) Cumulative probability plots of the mEPSC amplitude distribution. Averaged traces (**I**) of sEPSC (continuous line) and mEPSC (dotted line) and multiplicity index (**J**) for CA1 neurons of WT and KO mice.

**Table 1.** Intrinsic passive properties of hippocampal neurons.

|  | WT | KO |
| --- | --- | --- |
|  | (*n*=6) | (*n*= 8) |
| V<sub>m</sub> (mV) | -64.78±0.51 | -68.11±1.42 |
| R<sub>in</sub> (MΩ) | 149.05±3.40 | 170.15±16.90 |
| C<sub>m</sub> (pF) | 63.62±6.91 | 77.08±9.16 |
| τ | 9.29±0.87 | 12.92±1.65 |
| Number of AP | 9.67±1.72 | 10±0.92 |
| Firing frequency<sub>200pA</sub> | 11.5±4.97 | 12.65±0.96 |
| Firing frequency<sub>250pA</sub> | 13.83±4.66 | 14±1.82 |
| Firing frequency<sub>300pA</sub> | 15.4±5.12 | 14.5±2.63 |

Input resistance (R<sub>in</sub>) and resting membrane potential (V<sub>m</sub>) were measured in current clamp on the same cells used to construct I–V curves and determine threshold values. Whole-cell capacitance (C<sub>m</sub>) was measured in voltage clamp on cells used to measure ionic currents. No differences were significant.

**Table 2.** Morphological classification of synapses.

| Number of compound synapses | WT | KO |
| --- | --- | --- |
|  | (*n*=3) | (*n*= 3) |
| 1 contact | 80.9±23.34 | 208.0±17.65 |
| 2 contacts | 9.80±1.530 | 58.20±3.35 |
| 3 contacts | 0.80±0.200 | 4.40±0.8713 |
| Type of synapses |  |  |
|  | (*n*=3) | (*n*= 3) |
| Type I | 2.50±0.63 | 4.00±0.41 |
| Type II | 1.75±0.25 | 15.0±1.58 |
| Type III | 1.50±0.28 | 6.75±0.85 |
| Type IV | 0±0 | 2.25±0.47 |

'97 Data are mean ± SEM. \*Statistical significance determined by ANOVA ( $p < 0.05$ ) compared to WT '98 group. Type I = axon with two contacts; Type II= axon with one spine; Type III= axon with two '99 spines; Type IV= 2 axon with 2 spines. '00

To further examine differences between WT and KO, we examined the effect of PANX1 ablation on the Pr by estimating two principal forms of short-term synaptic plasticity: paired-pulse facilitation (PPF) ratio, measured as the relative strength of the second of two consecutive synaptic events, which is inversely related to Pr (*44*), and a use-dependent depression during a high-frequency stimulus train (*45*, *46*). Analysis of the PPF ratio revealed no changes between groups, suggesting no alteration in Pr evaluated by this mean (Fig. 3, A and B). Interestingly, we observed a remarkable difference in the EPSCs evoked by a train of 14 Hz (*47*, *48*) (Fig. 3, C to F). After the initial increase in the EPSC amplitude, in WT animals, we observed a progressive depression, consistent with a gradual depletion of the RRP recruited by the train (Fig. 3C). Interestingly, in KO neurons, we obtained persistent facilitation throughout the whole stimulatory train, which was reflected in the normalized curve of EPSCs significantly different from the WT group (Fig. 3D). Correspondingly, the cumulative current in KO neurons was considerably more extensive than WT neurons (Fig. 2E). Presynaptic changes also can be associated with modifications in the size of the readily releasable pool (RRP) of synaptic vesicles and the number of release sites or synaptic contacts (*46*). Thus, one approach to estimate the cumulative number of vesicles evoked by a train consists of plotting the cumulative peak of EPSCs divided by the mean amplitude of the mEPSC and fit over a linear region of the curve extrapolating back to zero (*45*). The linear back extrapolation revealed that the size of the RRP recruited by the train was significantly higher in KO neurons (Fig. 2F).

**Fig. 3.**
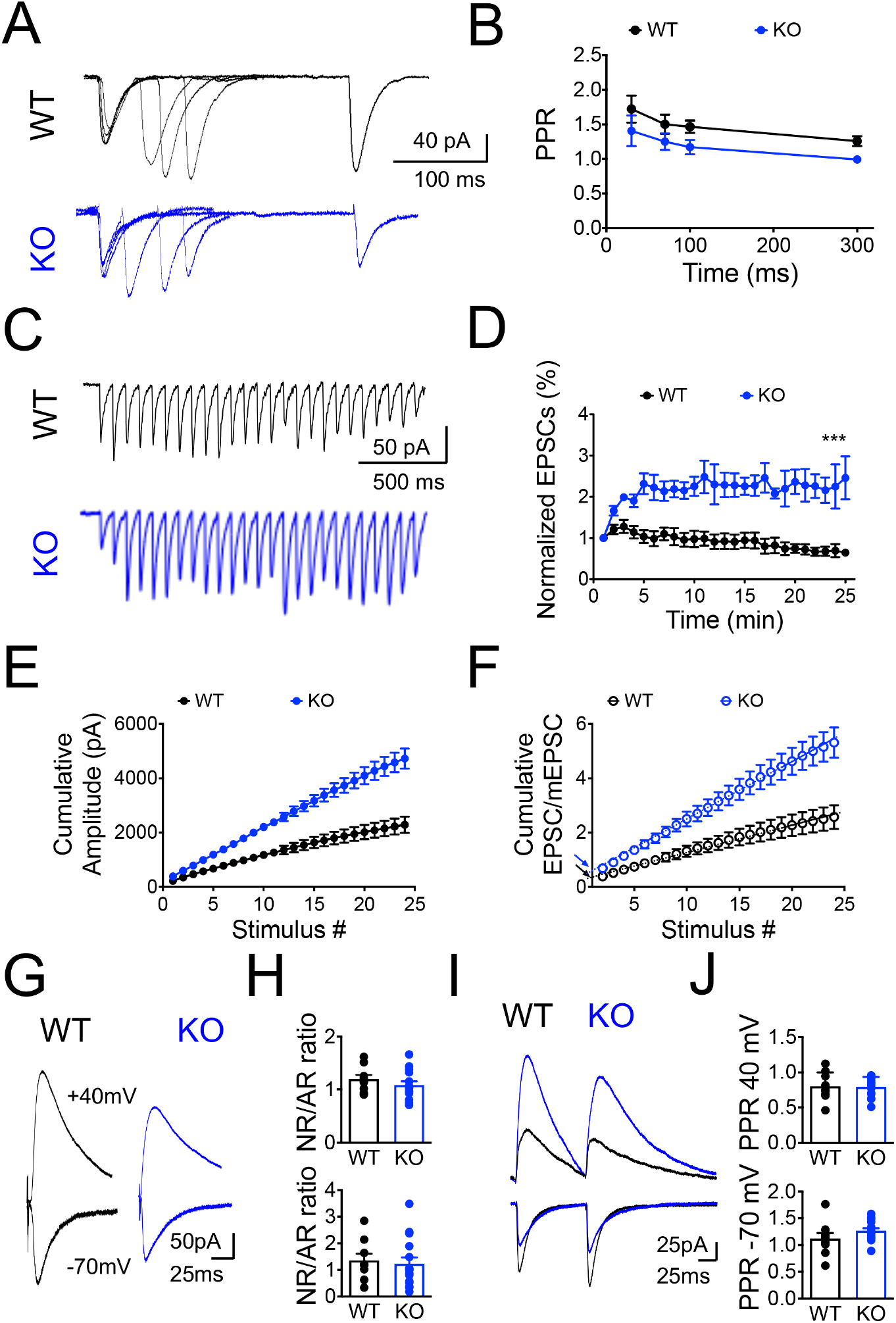
Readily releasable pool (RRP) and vesicle release probability are increased in PANX1 KO synapses. Representative traces (**A**) and analysis of paired-pulse facilitation ratio (**B**) in CA1 neurons of wild type (WT, black) and PANX1-KO (KO, blue) mice. Representative traces (**C**) and analysis of EPSCs evoked by a train of 25 pulses at 14Hz (**D**). Plot of the cumulative EPSCs versus number of stimulus (**E**) and cumulative EPSC/mEPSC amplitude to the estimation of the effective RRP size (RRPtrain) by back-extrapolation of the to the y-axis (**F**). Black and blue arrows indicate the RRP size for WT and KO respectively. Representative traces (**G**) and analysis of AMPAR to NMDAR (AR/NR) ratio (**H**) recorded at 40 mV (top) and - 70 mV (bottom). Representative traces (**I**) and analysis of paired-pulse ratio (PPR) obtained at 50-ms interpulse interval (**J**) and recorded at 40 mV (top) and −70 mV (bottom).

Short-term synaptic plasticity, particularly synaptic depression can also be contributed by postsynaptic mechanisms such as desensitization or saturation of AMPARs (*49*); we tested for a contribution of postsynaptic receptors in the increased facilitation observed in KO neurons by estimating an NMDAR- to AMPAR-mediated EPSCs ratio (NR/AR) and a paired-pulse ratio for AR-and NR-mediated EPSCs respectively (Fig. 3). We found that NR/AR was indistinguishable between groups, although NR- and AR-mediated currents were reduced in KO mice compared to that in WT littermates (Fig. 3, A and B). Similarly, we observed that the PPR of NR-and AR-mediated EPSCs were comparable between groups (Fig. 3, E and F).

These results indicate that spontaneous neurotransmission is unaltered in KO neurons but evoked glutamate release seems prompt to respond to activity demand likely by utilize different molecular machinery, segregated at distinct postsynaptic sites as has been previously reported (*50*). Furthermore, PANX1 ablation enhances neural excitability and suggest that KO neurons could hold homeostatic-like mechanisms to maintain synaptic transmission.

### Increased dendritic arborization and spine maturity of hippocampal CA1 pyramidal neurons from PANX1-KO mice

Since PANX1 ablation might induce adjustments in network connectivity and neuron excitability, we analyzed the impact of Panx1 ablation on the morphology of Golgi-stained neurons at dendrite and dendritic spine levels (Fig. 4). A significantly higher dendritic branching and dendritic length were observed in hippocampal neurons from KO mice compared to age-matched WT animals (Fig. 4, A to E). In addition, on average, KO neurons exhibited larger dendrites (Fig. 4B) with a higher total number of branch points (Fig. 4C) in both apical and basal dendrites (Fig. 4D), showing a higher number of dendrites throughout the proximal to distal distance from the soma in *stratum radiatum* layer (Fig. 4E). Furthermore, Sholl’s analysis also revealed that KO neurons displayed a significantly increased branch order indicative of greater dendritic complexity in the apical and basal compartments of the CA1 region (Fig. 4F). Concomitantly, analysis of dendritic segments of Golgi-impregnated neurons (Fig. 4G) revealed similar dendritic spines density (Fig. 4H), but a significantly increased dendritic spine length (Fig. 4I) in basal dendrites between wild type and KO animals. Moreover, KO neurons displayed a higher number of dendritic spines throughout the apical dendritic tree (Fig. 4J) that are also longer than WT neurons. Interestingly, when we classified dendritic spines based on their morphology in mature (mushroom- and cup-shaped) and immature spines (filopodia, thin and stubby). We observed that, in general, KO neurons exhibited a greater proportion of mature spines than WT neurons (Fig. 4, K to L) indicating that the absence of PANX1 also induces structural changes in spine morphology. Together these data suggest that the lack of PANX1 promotes a significant modification in neuronal morphology.

**Fig. 4.**
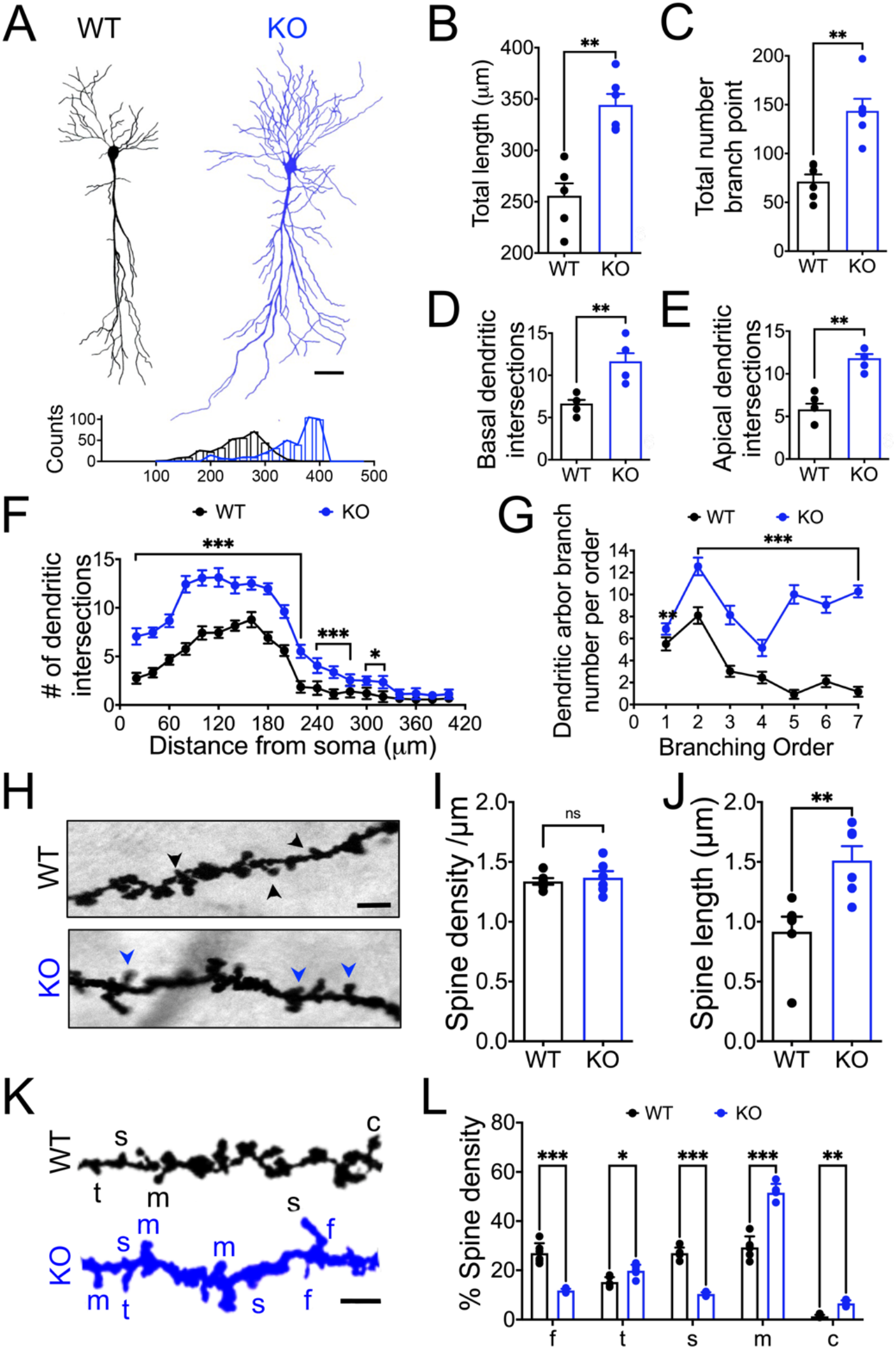
Enhanced dendritic arborization and spine maturation in PANX1KO neurons. (**A**) Representative drawings of Golgi stained CA1 neurons (top) and histogram distribution of total dendritic length (bottom). Sholl analysis of the averaged total dendritic length (**B**), dendritic branches (**C**), basal (**D**) and apical dendritic intersections (**E**) in CA1 neurons of wild type (WT, black) and PANX1-KO (KO, blue) mice. Number of intersections as a function of the distance from soma (**F**) and the branch order (**G**). Representative images of dendritic segments with dendritic spines (arrowheads) (**H**) and analysis of the spine density (**I**) and spine length (**J**). Pseudo-colored images of dendritic segments as in (**H**) showing different types of dendritic spines, filopodium (f), thin (t), short (s), and mushroom (m) types (**K**).

### Structural and molecular remodeling of synapses in hippocampal CA1 pyramidal neurons from PANX1-KO animals

Next, we examined the effect of PANX1 ablation on the synaptic structure by electron microscopy (Fig. 5). We found that CA1 hippocampal neurons from KO mice frequently exhibited compound synapses, either multi-innervated spines or an axon terminal contacting multiple spines (Fig. 5, A and B; Table 1; fig. S1). Furthermore, analysis of compound synapses revealed an increase in the proportion of multiple synaptic contacts in KO mice compared to WT animals (Fig. 5, A and B; Table 1; fig. S1). Moreover, KO neurons displayed a higher average number of synaptic contacts throughout the apical dendritic tree (Fig. 5C). According to larger and matured spines (Fig. 4, L and K), we found an increase in the PSD size (Fig. 5D). Remarkably, the number of small clear vesicles was significantly higher in KO spines compared to WT (Fig. 5, E and F), either total vesicle docked to the axon terminal membrane as well as docked vesicles at the active zone (Fig. 5, G to H).

**Fig. 5.**
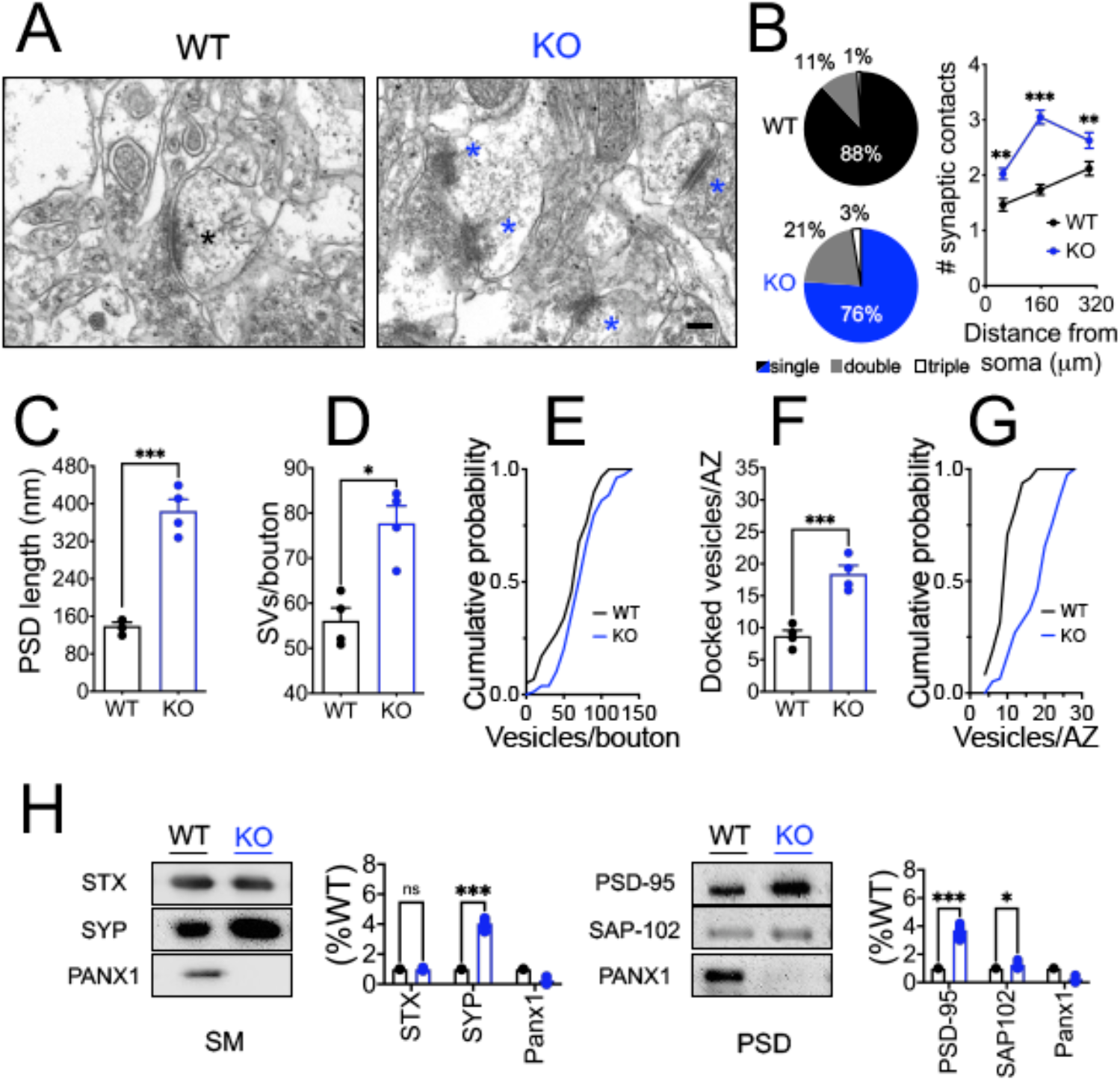
Multiple contacts, higher number of docked vesicles and PSD length in PANX1KO synapses. Representative transmission electron microscopy photographs of asymmetric synapses (**A**) and analysis of the percentage of contacts and number of contacts as a function of the distance from soma (**B**), PSD length (**C**) number of synaptic vesicles per bouton (**D**), cumulative probability of the docked vesicles distribution (**E**), docked vesicles at the active zone (**G**), and Ccumulative probability of the vesicles/AZ distribution in CA1 *Stratum radiatum* area of wild type (WT, black) and PANX1-KO (KO, blue) mice. Magnification 43000X, bar: 500nm. Representative blots and densitometric analysis of synaptic proteins levels in PSD-enriched and synaptic membranes avoided of PSD (SM)-enriched fractions (**H**).

We next tested whether the PANX1 ablation causes a major effect on synaptic protein content. Early studies have shown that PANX1 is expressed by neurons and glia (*51*) and accumulates preferentially in the postsynaptic density (PSD) of cortical and hippocampal neurons, colocalizing with postsynaptic proteins such as AMPAR and PSD-95 (*52*). Accordingly, we verified these observations by isolating hippocampal synaptosomes and subsequent separation into PSD-enriched fractions and synaptic membranes avoided of PSD (SM) (Fig. 5I). Interestingly, we detect the presence of PANX1 in both PSD-and SM-enriched fractions suggesting that Panx1 also could be present in the presynaptic compartment and could act on both levels. Furthermore, more recent works have reported a higher expression of postsynaptic protein such as PSD-95 and glutamate receptors in embryonic and adult PANX1 KO brains (*31*). Although no apparent change in the overall protein patterns were revealed by Coomasie blue-stained gel electrophoresis (fig. S2A), western blot from whole hippocampal tissue shows a modification in several synaptic proteins (fig. S2, C and D). We found a significant increase in presynaptic proteins such as synaptophysin (SYP) and postsynaptic proteins including, PSD-95 and SAP-102, in both whole hippocampal tissue (fig. S2, C and D) and synaptic enriched fractions (Fig. 5I). Overall, these results suggest that Panx1 lacking produces a major restructuring of the pre-and post-synaptic composition, consistent with the increase in synaptic connectivity.

### Lack of PANX1 promotes F-actin formation in hippocampal neurons via activation of Rac1 and depression of Rho A GTPases

Because the actin cytoskeleton dynamically regulates the morphology of dendrites and spines, the trafficking of glutamate receptors, and the molecular organization of PSD (*53*), we explored if the possible molecular mechanisms involved in the morphological changes observed in PANX1-KO neurons are related to cytoskeletal remodeling. Actin exists in a dynamic equilibrium between filamentous (F-actin) and monomeric (G-actin) forms, which are abundant in both presynaptic and postsynaptic compartments (*53*). We isolated actin filaments and monomers from hippocampal slices to estimate an F-actin/G-actin ratio (F/G) as a measure of actin polymerization. We found that the F/G ratio was significantly higher in KO than WT slices (Fig. 6A), suggesting that PANX1 ablation affects actin polymerization. Additionally, we investigated the impact of PANX1 ablation on F-actin content by visualization of Phalloidin fluorescence, a toxin that specifically binds to F-actin but not to G-actin (*54*). Accordingly, we observed that KO brain slices revealed higher phalloidin reactivity than WT mice (Fig. 6B). Quantification of the average fluorescence intensity of the dendritic layer of the CA1 region was significantly higher in KO slices (Fig. 6C), further indicating that PANX1 ablation promotes F-actin assembly. We reasoned that if PANX1 ablation promotes the formation of F-actin, then regulatory proteins that control actin dynamics and organization could mediate this effect. Thus, we evaluate the levels of actin-binding proteins (ABPs) and the expression of the Rho family GTPases, which are master regulators of the actin cytoskeleton (*55*). Western blot analysis revealed greater Arp3, Drebrin, Cortactin 1, Rac1, Cdc42, and RhoA levels in hippocampal homogenates from KO compared to WT samples (Fig. 6, D and E), indicating that the PANX1 deficiency affects actin remodeling by altering the expression of ABPs and Rho GTPases.

**Fig. 6.**
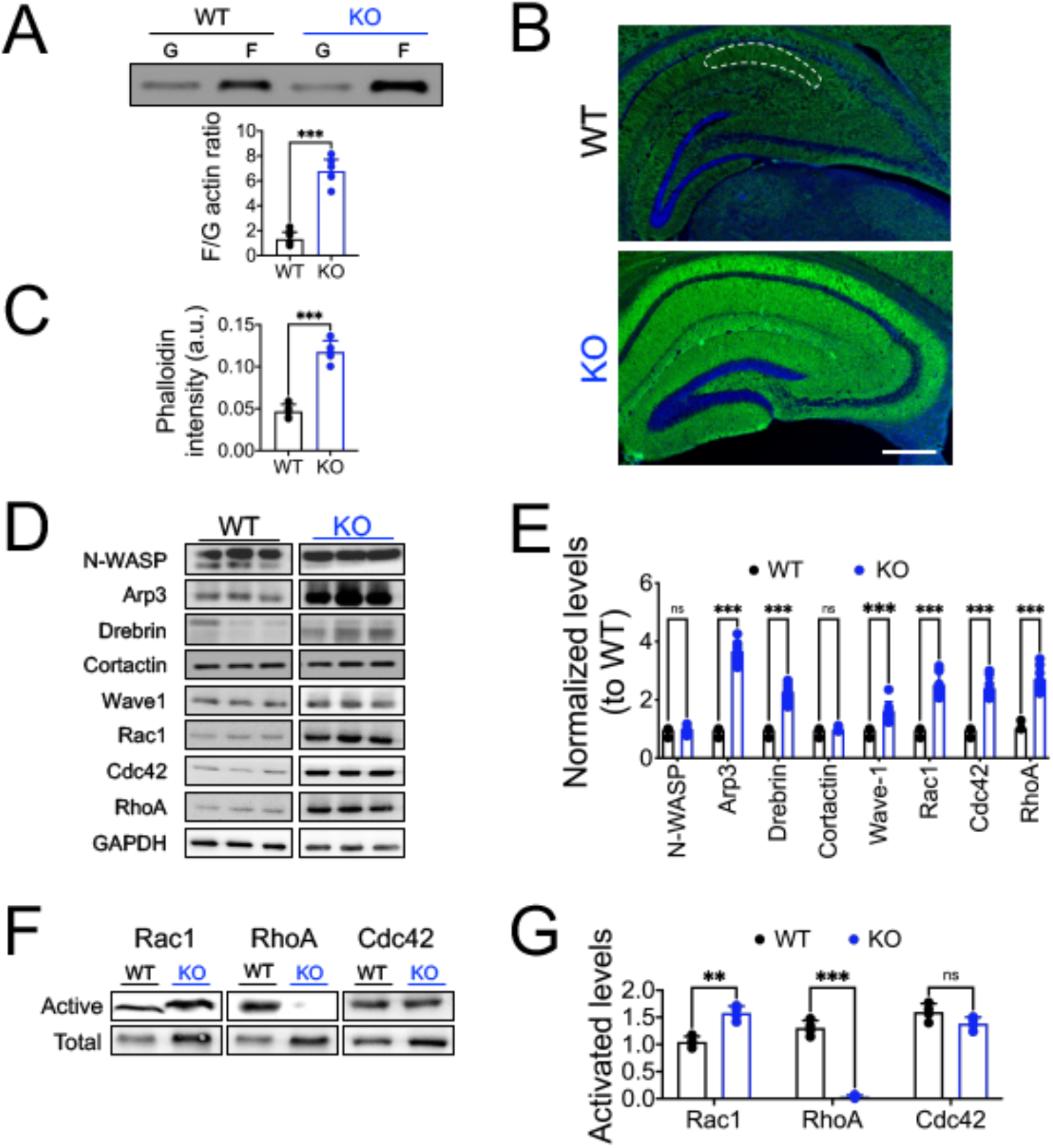
Increased actin polymerization and Rac1 activity in PANX1KO hippocampi. (**A**) Representative blots (top) and densitometric analysis of the relative number of monomers (G) and filaments (F) of actin (bottom), in hippocampal lysates of wild type (WT, black) and Panx1-KO (KO, blue) mice. (**B**) Representative micrographs showing Phalloidin-rhodamine staining of F-actin of CA1 region in hippocampus. Scale bar: 50 μm. (**C**) Quantification of phalloidin intensity. (**D**) Representative blots and densitometric analysis of small Rho GTPases and synaptic actin-binding proteins levels (**E**). (**F**) Representative blots and densitometric analysis of the relative ratios of the active Rac1, Cdc42 and RhoA proteins (**G**). All data are expressed as mean ± SEM of WT (n=5) and Panx1-KO (n=6), *p<0.005, **p<0.001.

Rho GTPases are important molecular switches transducing extracellular signals to the actin cytoskeleton (*53*, *55*). RhoA, Rac1, and Cdc42 are members of the Rho GTPases family implicated in the maintenance and reorganization of dendritic structures (*56*, *57*). Since the Rho GTPase family regulates actin dynamics, and PANX1 ablation seems to modulate actin polymerization and the expression of actin-related proteins, we hypothesize that PANX1 channels may control the steady-state of the activity levels of the Rho GTPases. To assay for activated GTPases, we employed GST fused to the GTPase-binding domain of PAK1, WASP, and Rhoketin, which bind to the active, GTP-bound forms of Rac1, Cdc42, and RhoA (*58*). We observed significantly increased levels of the active forms of Rac1 in KO mice compared to WT sample, and unexpectedly we also found almost absent levels of activated RhoA in KO tissue (Fig, 6 F and G). This result are consists with the antagonistic roles of Rac1 and RhoA in dendritic morphology (*59*). On the contrary, there were no significant differences in activated Cdc42 levels.

All these data support the hypothesis that PANX1 channels are associated with the actin cytoskeleton and suggest that modifications in the expression of PANX1 could directly impact on processes rely on the neuronal actin cytoskeleton.

## Discussion

PANX1 channels are non-selective channels essential for cellular communication under physiological and pathological conditions (*4*, *60*). In the CNS, they have been proposed to act as negative modulators since deletion, knockdown, or pharmacological inhibition of PANX1, enhance glutamatergic neurotransmission, LTP, and neurite outgrowth (*29*, *30*, *32*). On the other hand, changes in their expression are associated with neuronal hyperactivity and death, and aberrant Panx1 activity has been observed in several brain disorders, including ischemia (*23*, *24*), epilepsy (*25*, *26*) and Alzheimer’s disease (*28*). Therefore, changes in the expression and activity of PANX1 channels can be critical to the proper functioning of brain circuits and especially relevant for synaptic disorders. This study explores the consequences of long term PANX1 ablation in cellular and synaptic properties of adult hippocampal CA1 neurons. Using electrophysiological and structural studies, we support an important role of actin cytoskeleton orchestrating compensatory structural modifications in synapses without impact on its functionality.

### Long-term PANX1 ablation affects neuronal excitability but preserving spontaneous release

Under intense activity such as during the stimulation to evoke input/output curves, either the absence or the blockade of PANX1 induces an increase in synaptic responses (*29*, *30*). The mechanism proposed involves a reduction in the extracellular adenosine levels because of the lack of ATP release by PANX1 channels (*30*). However, it is still unknown how PANX1 could affect synaptic transmission under basal conditions. Therefore, we monitored glutamatergic neurotransmission and synaptic connectivity to determine whether long-term PANX1 deficiency elicits adaptations to regulate synaptic strength. As a result, we found that CA1 pyramidal neurons from KO mice exhibited a lower threshold for action potential discharge and fired more action potentials in response to a current ramp. Nevertheless, we found that PANX1 ablation does not affect the intrinsic membrane properties of CA1 neurons, neither produces apparent difference in I-V curves were observed, although we cannot rule out that selective ionic currents might be affected.

In this regard, similar changes in the threshold for AP discharges have been associated with neuronal maturation and homeostatic mechanisms (*61*, *62*). Several studies have shown that changes in selective ion channels affect the intrinsic excitability of hippocampal neurons (*63*). For instance, potassium currents I_D_ (*64*) and I_h_ (*65*) contribute to the homeostatic modifications of intrinsic excitability in CA1 pyramidal neurons. Similarly, a persistent TTX-sensitive sodium current (I_NAP_) reduces the spike threshold and triggers repetitive firing under muscarinic activation in CA1 neurons (*66*). On the other hand, changes in the KCC2 transporter, which regulates neuronal chloride gradients and GABA signaling, also modulates intrinsic neuronal excitability (*67*), suggesting that GABA transmission can also be an important factor affected in KO neurons. Thus, although the evaluation of inhibitory transmission has not been accomplished in the present study, modifications in the E/I balance would be an aspect to consider ahead and require further investigation.

In addition to a large pore conformation with high conductance (100-550 pS), mediating a non-selective ionic flux and ATP release (*6*, *14*, *16*), PANX1 channels also show a constitutive small pore activity characterized by low conductance (50-80 pS) driving a chloride permeability at negative voltages and outwardly rectifying current-voltage relations (*7*–*10*), which could influence the electrical properties of the cells and can to explain the modifications in the action potential threshold upon PANX1 ablation. Accordingly, a recent report revealed a higher excitability in the hippocampus of PANX1-KO mice(*68*). CA1 pyramidal neurons exhibited a more positive resting membrane potential, a lower excitatory threshold, faster action potentials and a higher firing frequency compared to control littermates.

To test if excitability changes are reflected in synaptic transmission modifications, we record evoked (fEPSCs) and spontaneous (sEPSCs, and mEPSPs) synaptic events. We did not find significant differences either in the initial slope of fEPSPs or afferent fiber volley (FV) amplitude, which suggest non apparent change in efficacy of evoked excitatory synaptic transmission between KO and WT. However, AR-and NR-FP revealed that KO neurons exhibit an enhanced tendency to fire pop spikes consistent with an impairment in excitatory/inhibitory balance or a modification in the intrinsic excitability of the CA1 neurons. Since GABAergic neurons regulate the activity of principal neurons and modulate the oscillatory activity of neural networks, it should be further addressed inhibitory GABAergic transmission and oscillatory activity.

The latent higher excitability revealed by the changes in the firing threshold and the generation of pop-spikes seem not due to an increase in spontaneous basal synaptic transmission as the frequency and the amplitude of sEPSCs were indistinguishable between groups. Likewise, the frequency of mEPSCs were similar, indicating that spontaneous synaptic activity is normal in KO neurons (Fig. 1, D to F). However, the amplitude of mEPSCs was higher which could correlate with the higher excitability showed by KO neurons. These spontaneous events independent of action potentials inform about the potential locus for synaptic modification. The amplitude of mEPSCs is thought to reflect changes in the expression (number) or activity (conductance) of neurotransmitter receptors; meanwhile, changes in the frequency of mEPSCs has been related to presynaptic modifications such as the probability of neurotransmitter release (Pr) and/or the number of synapses (*69*). The fact that the amplitude of mEPSCs were higher in KO neurons suggests for a postsynaptic modification, although we cannot discard that lack of PANX1 affects the pre-synaptic transmission mechanisms.

Our observations differ from those reported upon PANX1 blockade or interfering with NMDAR-PANX1-Src kinase pathway induces an increase in the sEPSC frequency (*70*). Although Bialecki et al. described similar results in conditional KO neurons, those effects were seen in younger animals. Thus, when we evaluated synaptic responses in our conditions, we believed that long-term homeostatic mechanisms could occur that permit support synaptic transmission in the adult hippocampus without alterations in spontaneous neurotransmitter release. However, we cannot discard the contribution of presynaptic TRPV1-mediated facilitation of glutamate release upon PANX1 targeting in our experimental conditions.

### Increased size of ready releasable pool RRP of vesicles but normal release probability

We previously reported in the adult PANX1 KO an increased induction of LTP and a lower capacity to induce LTD, which was ever a potentiation instead a depression using standard electrophysiological protocols. (*29*, *30*). Despite that, chemical induction of LTD revealed that KO synapses indeed present the protein machinery to support this plastic modulation (*29*, *30*). Our present findings show that in response to a use-dependent depression during a high-frequency stimulus train, KO neurons exhibit a sustained response indicating a greater size of RRP of vesicles. This observation can explain the potentiated response observed in KO slices after the application of a protocol that induce an LTD in control animals(*29*, *71*).

An increase in glutamate release could result from increasing the number of Ca^2+^-responsive vesicles (i.e., an increase in the RRP), or an increase in the release probability (Pr). The examination of the paired-pulse facilitation (PPF) revealed no change between groups. Consistently with that, the use-dependent depression induces by a 14 Hz stimulus train revealed a reluctance of KO synapse to deplete the RRP vesicles evoked by the stimulus train. Consistently, the RRP size derived from the linear back extrapolation of the cumulative EPSCs divided by the mEPSC mean, was significantly increased at KO synapses, suggesting that the pool of vesicles immediately available for release could be greater in KO neurons.

Interestingly, recent evidence highlights the role of PANX1 channels in the homeostatic adjustment of synaptic strength in hippocampal cultures upon chronic inactivity (*72*). Using glutamate transporter 1 (vGlut1) immunodetection as an index of presynaptic strength, Rafael et al., reported that PANX1 channels are required for the compensatory vGlut1 upregulation in presynaptic terminals and the adjustment of synapse density under chronic inactivity.

### Long-term PANX1 ablation affects structural but not functional connectivity

The present study also showed that the multiplicity index, an estimation of the number of releasing sites or likely synaptic contacts, was significantly lower in KO neurons. This is in contrast with the ultrastructural findings that revealed a greater number of contacts in KO neurons. In this regard, the highly branched dendritic tree of central neurons defines how the cell can receive synaptic inputs from other neurons and strongly influences how these inputs are integrated to allow signal transmission and computation. Our results revealed a more complex morphology of CA1 neurons from KO neurons as revealed by the intersection profiles obtained by counting the number of dendritic branches at a given distance from the soma. KO neurons displayed longer dendrites with more ramifications in both apical and basal segments and higher branch order. Indeed, this greater dendritic complexity supposed that KO neurons would possess the morphology, structure, and protein composition for more increased connectivity. KO neurons exhibit a more significant percentage of mature-like dendritic spines, a higher proportion of multiple synaptic contacts, spines with a larger PSD size, and a higher number of synaptic vesicles per bouton or active zone. Accordingly, the results obtained under depletion experiments indicate that KO mice exhibit a greater size of the RRP as evidenced by the increase in the synaptic responses after the subsequent stimuli. Moreover, PSD-and SM-enriched fractions obtained from hippocampal synaptosomes disclose an enhanced expression of pre-and post-synaptic proteins suggesting that PANX1 ablation might induce compensatory modifications in both compartments. These data demonstrated that morphological changes in dendritic arborization and spines, along with a change in the synaptic protein composition, likely reflect a modification in the remodeling of the neuronal cytoskeleton. We tested this hypothesis by evaluating the organization of the actin-dependent cytoskeleton. Actin cytoskeleton transits in a dynamic equilibrium between two states, F-actin and G-actin forms, abundant in presynaptic terminals and postsynaptic dendritic spines (*53*). Furthermore, this equilibrium between G-actin and F-actin is finely and rapidly regulated during activity by many postsynaptic ABPs. In this study, we reported that KO neurons exhibit higher content of F-actin, suggesting that PANX1 ablation promotes a disequilibrium in actin dynamics towards polymerization. In support of a regulatory role of PANX1 over actin cytoskeleton, it has been previously reported a direct interaction between PANX1 and F-actin (*73*) and the actin-related protein Arp3 (*32*), which is the main component of the Arp2/3 complex involved in *de novo* nucleation and branching of F-actin (*74*). Moreover, PANX1 has been implicated in several cell behaviors reliant on actin rearrangements, down-regulation of its activity promotes neurite outgrowth, cell migration (*32*, *75*), and dendritic spines development (*31*), suggesting that PANX1 activity limits neuronal cytoskeleton remodeling. The mammalian Rho family of small GTPases are key regulators that control the organization and dynamics of the actin cytoskeleton (*55*). Among them, Rac1, Cdc42, and RhoA play a major role in spine dynamics, connecting signals from the postsynaptic neurotransmitter receptors to changes in ABPs and hence, actin polymerization/depolymerization (*56*, *57*). While Rac1 and Cdc42 activation promote spine formation and stabilization, RhoA activation leads to spine retraction and pruning [62, 65]. Accordingly, we also found that hippocampal tissue from KO mice exhibits enhanced Rac1, Cdc42, and RhoA levels along with selective ABPs including Arp3, Drebrin, and Wave-1 of actin remodeling. Interestingly, the active form of Rac1, but not Cdc42, was significantly increased in hippocampal homogenates from KO mice. In contrast, the activated form of RhoA was almost absent, consistent with their antagonistic roles in dendritic and spine morphology (*59*). Thus, it appears that PANX1 channels may control the steady-state activity levels of Rho GTPases Rac1 and RhoA, promoting the actin cytoskeleton dynamics to support morphological changes of synaptic structure.

These findings revealed a disbalanced equilibrium towards spines more mature supported by increased actin polymerization which implies a more static than plastic synaptic architecture. These observations also can explain the previously reported difficulty to induce LTD mechanisms which implies a retraction or reduction in the size and number of dendritic spines and a higher depolymerization of actin filaments. In fact, despite PANX1 ablation has been associated with enhanced glutamatergic neurotransmission, LTP, and neurite outgrowth (*29*, *30*, *32*), alterations in learning flexibility, such as novel object recognition, spatial memory, and anxious behaviors have been described in PANX1 KO mice (*30*, *71*). Nevertheless, the animals preserve their ability to learn, so compensatory or homeostatic mechanisms must operate to maintain the plasticity of the adult brain at cognitively significant levels. Therefore, changes in Panx1 expression and activity in the brain can be critical to the proper functioning of neural circuits and especially relevant for synaptic diseases.

Overall, the results presented here show that long term ablation of PANX1 channels promotes a sequence of functional and morphological changes in hippocampal CA1 neurons which can be critical to the proper functioning of neural circuits. Interestingly, our findings prove that these modifications rely on the Rho GTPase-dependent regulation of the actin cytoskeleton. Thus, the functional interaction between PANX1 channels and Rho GTPases could be of pivotal relevance considering that numerous brain disorders such as neuropsychiatric and neurodegenerative conditions develop with abnormalities in neuronal cytoskeleton.

## Materials and Methods

### Animals

The experiments were carried out in adult male (6-9 months old) C57BL/6 or PANX1-KO mice. The generation of PANX1-KO mice has been described previously (*76*). Mice were housed at 21 ± 1°C at constant humidity (55%) and in a 12/12 h dark-light cycle, with a light phase from 08:00 to 20:00. Food and water were provided *ad libitum*. All animal experiments were approved by the Ethical and Animal Care Committee of the Universidad de Valparaiso (BEA064-2015).

### Electrophysiology

Animals were deeply anesthetized with isoflurane (forane, B506, abbvie), the brains were quickly removed and submerged in a cold (4°C) dissection buffer (in mM: 124 sucrose, 2.69 KCl, 1.25 KH_2_PO_4_, 10 MgSO_4_, 26 NaHCO_3_, 10 glucose). Then, coronal hippocampal slices (350 μm) were cut with a Vibratome (WPI Instruments, model NVSLM1, FL, USA) and maintained for > 1 h at room temperature (20°C) in artificial cerebrospinal fluid (ACSF) composed, in mM: 124 NaCl, 2.69 KCl, 1.25 KH_2_PO_4_, 2 MgSO_4_, 26 NaHCO_3_, 2 CaCl_2_, and 10 glucose. The pH of the dissection buffer and ACSF were adjusted at 7.40 with NaOH or HCl and stabilized by bubbling carbogen (95% O_2_, 5% CO_2_). To perform electrophysiological recordings, the slices were transferred into a 2 mL chamber fixed to an upright Nikon Eclipse FN1 microscope stage (Nikon Instruments, TYO, JP) equipped with infrared differential interference contrast (DIC) video microscopy and a 40x water immersion objective. The slices were continuously perfused with carbogen bubbled ACSF (2 mL/min) and maintained at room temperature (22-24°C). Picrotoxin (PTX; 10 mM, Tocris) and tetrodotoxin (TTX; 0.5 mM) were added to the ACSF as needed. All chemicals were purchased from Sigma-Aldrich Chemistry (St. Louis, MO, USA), Tocris (Bioscience, Pittsburgh, PA, USA), and Cayman Chemical (Sarasota, FL, USA). Whole-cell recordings were performed from the soma of pyramidal neurons from the CA1 area of vHPC with patch pipettes (4–8 MΩ) filled with an internal solution containing, in mM: 100 Cs-Gluconate, 10 HEPES, 10 EGTA, 4 Na_2_-ATP, 10 TEA-Cl, and 1 MgCl_2_-6H_2_O, buffered to pH 7.2–7.3 with CsOH. Recordings were performed in a voltage-clamp configuration using an EPC-7 patch-clamp amplifier (HEKA, Instruments, MA, USA). The holding potential (V_h_) was adjusted to 0 mV to record excitatory postsynaptic currents (EPSCs). In the voltage-clamp configuration, the series resistance was compensated to ~70%, and neurons were accepted only when the seal resistance was > 1 GΩ, and the series resistance (7–14 MΩ) did not change by > 20% during the experiment. The liquid junction potential was measured (~6mV) but not corrected. Voltage-clamp data were low-pass filtered at 3.0 kHz and sampled at rates between 6.0 and 10.0 kHz using an A/D converter (ITC-16, InstruTech, MA, USA) and stored with the Pulse FIT software (HEKA, Instruments, MA, USA). The Pulse Fit program was used to generate stimulus timing signals and transmembrane current pulses. The recording analysis was made offline with pClamp software (Clamp-fit, Molecular Devices, CA, USA). eIPSCs were evoked by stimulation and recording in *Stratum pyramidale*. Stimulation was made with a concentric bipolar electrode [60 mm diameter, tip separation ~100 mm (FHC Inc., ME, USA)]. Averages of EPSCs were obtained by repeated stimulation at 0.3 Hz. Paired pulse facilitation and use-dependent depression. Paired pulse stimulus was applied at four different intervals (30, 70, 100, and 300 ms) and was calculated as R2/R1, where R1 and R2 correspond to peak amplitudes of the first and second eEPSCs, respectively. Use-dependent synaptic depression was analyzed using 15 Hz bursts of 25 stimuli every 60 s (~3 V, 200 ms). Six to ten responses at each intensity were averaged to compute the EPSC amplitude. sEPSC recordings were continuously recorded at 0mV for 30 min, under PTX. In addition, miniature EPSCs (mEPSCs) were recorded after adding the voltage-gated sodium channel blocker, TTX (0.5 mM), to the bath. To estimate the size of the readily releasable pool of vesicles, we use an approximation as previously described (*45*, *47*, *77*). The amplitudes of EPSCs evoked by a high-frequency train (25 pulses at 14 Hz) were measured and summed, and a graph was made of the cumulative EPSC. Then these values were divided by the amplitude of mEPSCs, as they represent the cumulative number of vesicles (*45*). The readily releasable pool liberated by the stimulus train (RRPtrain) was then determined by fitting over a linear region of this curve and extrapolating back to the y-axis (a straight line was fitted to the final 5 points of the cumulative EPSC). The y-intercept corresponds to RRPtrain. Furthermore, the probability of release related to the train (Ptrain) was estimated by dividing the amplitude of the first EPSC by RRPtrain (*47*, *77*). Finally, multiplicity was estimated as previously described (*42*, *43*). Multiplicity index was calculated as the mean amplitude of action potential-driven events (a) divided by mean quantal size (q: mean amplitude of mEPSC recorded in TTX). The a values were determined for each cell, subtracting the contribution of mEPSC to the pool of events collected in the absence of TTX, using the expression for a:

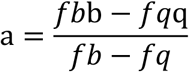

Where *f_b_* and *f_q_* denote the mean frequency values from events recorded before and after the addition of TTX to the perfusion media, respectively, and b is the mean amplitude of both action-potential-driven sEPSC and mEPSC.

For extracellular field recordings, transverse hippocampal slices (350-μm-thick) were dissected in ice-cold (4 °C) dissection buffer, in mM: 212.7 sucrose, 2.6 KCl, 1.23 NaH_2_PO_4_, 26 NaHCO_3_, 10 dextrose, 3 MgCl_2_, and 1 CaCl_2_, using a vibratome (Leica VT1200S, Leica Microsystems, Nussloch, GE). Slices then were recovered for 1 h at room temperature in ACSF, in mM: 124 NaCl, 5 KCl, 1.25 NaH_2_PO_4_, 26 NaHCO_3_, 10 glucose, 1.5 MgCl_2_, and 2.5 CaCl_2_. Both dissection buffer and ASCF were pH adjusted at 7.40 and bubbled with carbogen (95% O_2_, 5% CO_2_). After recovery, the slices were placed in a submersion recording chamber perfused with ACSF (28 ± 0.5 °C; 2 mL/min). Field excitatory postsynaptic potentials (FPs), basically AMPAR-mediated FPs (AR-FPs), were evoked by stimulating Schaffer collaterals with 0.2 ms pulses delivered through concentric bipolar stimulating electrodes (FHC Inc., ME, United States) and recorded in CA1 *Stratum radiatum* using glass electrodes filled with ACSF (1 MΩ). FPs were amplified, low-pass filtered (1700 Differential AC Amplifier, A-M Systems), and then digitized (NI PCI-6221; National Instruments) for measurement. Baseline responses were recorded at 0.033 Hz. Next, NMDAR EPSPs (NR-FP) were isolated by applying CNQX (10 mM) in ACSF containing 2 mM calcium and 0.1 mM magnesium. After 30 min of CNQX pre-incubation, NR-FPs were recorded. Finally, basal synaptic transmission was assayed by determining input-output relationships from FPs generated by gradually increasing the stimulus intensity; the input was the peak amplitude of the fiber volley (FV), and the output was the initial slope of FP. Data were monitored, analyzed online, and reanalyzed offline using a homemade program based on Igor software (Wavemetrics, Lake Oswego, OR, USA). Representative traces are an average of four consecutive responses.

### Golgi staining to dendritic morphology visualization

Neuron morphology and dendritic spines were measured using FD Rapid GolgiStain Kit according to the manufacturer’s guidelines (FD NeuroTechnologies, Columbia, MD, USA). Briefly, dissected mouse brains were immersed in Solution A/B for two weeks at room temperature in dark conditions. Next, brains were placed in Solution C for 24 h in the dark. Afterward, coronal slices 150 μm thick were obtained using a semi-automatic cryostat microtome (Kedee KD-2950, Jinhua, ZJ, CN) at - 20°C and mounted on gelatin-coated microscope slides with Solution C. The sections were allowed to dry naturally at room temperature, then placed in a mixture of Solution D/E for 10 min. Next, sections were rinsed twice in Milli-Q water for 4 min each time. Finally, the sections were dehydrated, cleared in xylene, and mounted using Permount mounting media (Sigma-Aldrich, Burlington, MA, USA).

### Morphometric analysis

For morphometric analysis, digital images were taken of individual well-impregnated pyramidal neurons of the CA1 region using a Leica DM500 microscope (Leica Microsystems Inc., Buffalo Grove, IL, USA) equipped with a 40X objective, and ICC50W digital camera (Leica Microsystems Inc., Buffalo Grove, IL, USA). Neurons were then drawn with the aid of a camera lucida (Leica drawing device L3/20) attached to the microscope (Leica Microsystems Inc., Buffalo Grove, IL, USA) and digitalized in 1.200 × 1.200 dpi resolution to morphometric Sholl analysis using the Neuroanatomy and Simple neurite tracer plugins of Fiji software. In short, a series of concentric spheres (centered around the soma) were drawn with an intersection interval of 20 μm to calculate the number of dendrites crossing each sphere. This analysis was done for basal and apical dendrites and was plotted against the distance from the soma. Three parameters were used to determine dendritic morphology and complexity: 1) the number of dendrites; 2) the total dendritic length, including all dendritic branches; 3) the number of dendritic branches; and 4) the branch order. All morphological analysis was performed blind to experimental conditions.

For spines analysis, dendritic segments from primary and secondary branches of 10 to 20 μm in length were selected randomly. Images were acquired using a Leica DM500 microscope (Leica Microsystems Inc., Buffalo Grove, IL, USA) equipped with a 63X oil HCPL APO objective (NA 1.40) and ICC50w digital camera (Leica Microsystems Inc., Buffalo Grove, IL, USA). Spine density was defined as the number of spines per 10 μm. Dendritic spines were classified according the following parameters(*78*): Long, thin without-head protrusions of more than 2 um long were classified as immature filopodia; wide-head (>0.6 um width) and short protrusions (<1 um) were classified as mature mushroom-spines; wide-head/without neck protrusions (length: width ratio <1 um) were classified as stubby-spines; thin and short (<2 um) headed protrusions were classified as thin-spines. Cup-shaped protrussions were classified as branched spines. The spines parameters were analyzed using ImageJ (version 1.49v; NIH, Maryland, USA).

#### Electron Microscopy on hippocampal CA1 region

Electron microscopy analysis was processed for histological studies as previously reported (*71*). Mice were transcardially perfused with a mixture of 4% PFA and 0.5% glutaraldehyde, followed by post-fixation in the same mixture ON at 4°C. Brain tissue blocks were trimmed in the CA1 area of the hippocampus and dehydrated in a graded series of ethanol, infiltrated in 1:1 volumes of 100% ethanol and 100% LR White (EMS) during 4 h, immersed in 1.5% OsO_4_ in 0.1 M sodium phosphate buffer (pH 7.4) for 2 h and then embedded in epoxy resin which was polymerized at 50°C ON. Ultrathin sections (90 nm) were made using an ultramicrotome (Leica Ultracut R, Leica Microsystems, Nussloch, Germany) and contrasted with 1% uranyl acetate and lead citrate and located on nickel 300 mesh grids (Ted Pella Inc., Redding, CA, USA). Grids were observed under a transmission electron microscope Philips Tecnai 12 operated at 80 kV (FEI/Philips Electron Optics, Eindhoven) equipped with a digital micrograph camera (Megaview G2, Olympus). Synapses from 1 littermate pair have been analyzed by an experimenter who was blind to the genotype, and the following parameters were analyzed: total number of synaptic vesicles, number of docked vesicles, number of vesicles within the active zone, PSD length, and number of synaptic contacts.

### Subcellular fractionation-Synaptosome isolation

Synaptosomes were extracted from the hippocampus of 6 m.o adult male mice as we previously reported (*71*). Hippocampi were homogenized using a Dounce Tissue Grinder in ice-cold homogenization buffer containing, in mM: 320 sucrose, 4 HEPES, 1 EGTA, and protease and phosphatase inhibitor’s cocktail buffered to pH 7.4. The homogenate was centrifuged at 1,800 rpm for 10 min at 4°C (Beckman F0630 rotor) obtaining a supernatant (S1) which was collected whereas the pellet (P1) was discarded. Then, S1 was centrifuged at 30,000 rpm for 30 min at 4°C (Beckman S4180 rotor). The obtained pellet (P2) containing the membrane proteins was re-suspended in homogenization buffer, layered on the top of a discontinuous sucrose density gradient (0.32/1.0/1.2 M) and subjected to ultracentrifugation at 86,000 rpm (Beckman SW-60ti rotor) for 2 h at 4°C. Afterwards, both the sediment and sucrose 0.32/1M interface were discarded, whereas material accumulated at the interface of 1.0 M and 1.2 M sucrose containing synaptosome fraction was collected (SP1). SP1 was diluted with lysis buffer to restore the sucrose concentration back to 320 mM and remained on ice with gently agitation for 30 min. Then, SP1 was centrifuged at 30,000 rpm for 30 min. The pellet obtained (PS1) was resuspended in a gradient loading buffer, loaded on 0.32/1.0/1.2 M discontinuous gradient, and centrifuged at 86,000 rpm for 2 h. The sucrose 1/1.2M interphase, synaptosome fraction 2 (SP2), was recovered and delipidated in a delipidating buffer. Next, SP2 was diluted with a filling buffer to restore the sucrose concentration and then centrifuged at 30,000 rpm for 1 h. The sediment obtained (PS2) was washed with 50 mM HEPES-Na and centrifuged at 86,000 rpm for 20 min. The final sediment obtained (PS3), containing post-synaptic densities (PSD), was re-suspended in 50 mM HEPES-Na and homogenized. PS2 or PSD fractions were quantified for protein concentration using the Qubit^®^ Protein Assay Kit (Thermo Scientific Rockford, IL, USA).

### Immunoblotting

Total proteins and synaptosomal fractions were processed for western blotting as previously described (*79*). Samples for total tissue proteins were homogenized in ice-cold lysis buffer containing: 150 mM NaCl, 10 mM Tris-Cl, pH 7.4, EDTA 2 mM, 1% Triton X-100 and 0.1% SDS, supplemented with a protease and phosphatase inhibitor cocktail (Thermo Scientific, Rockford, IL, USA) using a Potter-homogenizator. Protein samples were centrifuged twice for 5 min at 14,000 rpm. (4°C). Protein concentration was determined using the Qubit^®^ Protein Assay Kit (Thermo Scientific Rockford, IL, USA). For both cases, 40 μg of protein per lane were resolved by 10% SDS-PAGE, and 12% SDS-PAGE for actin-proteins followed by immunoblotting on PVDF membranes (BioRad, California, USA) and probe with specific antibodies against Panx1 (rabbit anti-Panx1, ABN242 Merck; 1:1000), PSD95 (mouse anti-PSD95, MAB1596 Merck; 1:1000), SAP102 (mouse anti-SAP102, NeuroMAP; 1:500), Synaptophysin (goat anti-SYP, sc-9116 Santa Cruz; 1:2000), Syntaxin (rabbit anti-STX, sc-5899 Santa Cruz; 1:1000), Arp3 (rabbit anti-Arp3, 07272 Merck; 1:1000), N-WASP (mouse anti-N WASP, Santa Cruz; 1:2000), Rac1 (mouse anti-Rac1, sc-217 Santa Cruz; 1:1000), RhoA (mouse anti-RhoA, sc-418 Santa Cruz; 1:1000), Cdc42 (mouse anti-Cdc42, sc-8401 Santa Cruz; 1:1000) and GAPDH (mouse anti-GAPDH, sc-47724, Santa Cruz; 1:1000). After primary antibody incubation and washing incubation with a secondary anti-mouse HRP antibody (1:5000), anti-rabbit HRP antibody (1:5000) or with anti-goat HRP antibody (1:5000) was performed for 1 h and ECL (Pierce, Thermo Scientific, Rockford, IL, USA) visualized detection. Immunoreactive bands were scanned and densitometrically quantified using Image J software (version 1.49v; NIH, Bethesda, MD, USA). Total and synaptosomal fractions proteins data were normalized to GAPDH and expressed as a % of control group (wild type, WT).

### Rho GTPases activation assay

Hippocampal slices were stabilized in a chamber with oxygenated (95% O_2_ and 5% CO_2_) ACSF, pH 7.4 for 1 h. The GTP loading of Rac1, Cdc42, and RhoA was measured using a Rho GTPase Activation Assay Combo Biochem Kit (BK030, Cytoskeleton, USA), according to the manufacturer’s recommendations. Briefly, the hippocampal lysates were incubated at 4°C on a rocked for 1 h with Rac/Cdc42 (PAK1 PAK-binding domain) or RhoA (Rhoketin-binding domain) beads which binds specifically to GTP-bound, and not GDP-bound. Agarose beads were collected by centrifugation (for 3 min at 5,000 g at 4°C), washed and the immunoprecipitated resolved on 12% SDS-PAGE and detected via Western blot analysis using Cdc42 (mouse anti-Cdc42, ACD03 Cytoskeleton; 1:250), Rac1 (mouse anti-Rac1, ARC03 Cytoskeleton; 1:500) and RhoA (mouse anti-RhoA, ARH05 Cytoskeleton; 1:500), and visualized by ECL (Pierce, Thermo Scientific, Rockford, IL, USA). Rac/Cdc42 or RhoA activation values were expressed as the ratio of Rac1-GTP, Cdc42-GTP, or RhoA-GTP against the total of proteins of each Rho GTPases in the crude extract.

### F-actin quantification

Hippocampal slices were fixed at room temperature with 4% paraformaldehyde and 15% sucrose for 30 min and maintained in PBS buffer to evaluate total F-actin. After that, slices were cut into 25 um sections using a cryostat (Leica CM1900) and incubated with 1 μM of the F-actin-binding toxin phalloidin-tetramethyl-rhodamine-B (Sigma Aldrich, P1951) for 1 h. Additionally, hippocampal sections were co-stained with DAPI (Sigma Aldrich; 1:1000) to label nuclei. Images were acquired in a confocal microscope (upright Eclipse Nikon 80i), and the fluorescence intensity was measured using the Image-J software (version 1.49v; NIH, Bethesda, MD, USA). All the image preparations were performed by Photoshop software (Adobe, California).

### F-actin/G-actin assay

The filamentous (F-actin) and monomer actin (G-actin) content were quantified as previously described (*80*). Briefly, hippocampal slices previously stabilized for 1 h with oxygenated ACSF were lysed and homogenized in the presence of conditions that stabilize F-actin and G-actin using an F/G actin commercial assay (F/G actin *in vivo* assay BK037, Cytoskeleton Inc., USA). Then, the homogenates extracts were ultracentrifuged at 100,000 g for 1 h at 37 °C to separate F-actin (pellet) and G-actin (supernatant) fractions. F-actin pellet was resuspended in a depolymerizing buffer (BK037, Cytoskeleton Inc., USA) and then F- and G-actin fractions were diluted in loading buffer (50 mM Tris–HCl, 2% SDS, 10% glycerol, 1% beta-mercaptoethanol and bromophenol blue). The samples were resolved on 12% SDS-PAGE and detected via Western blot analysis using actin (rabbit anti-actin, BK037 Cytoskeleton Inc.; 1:500) and visualized by ECL (Pierce, Thermo Scientific, Rockford, IL, USA). The ratios of F-actin or G-actin in hippocampal slices were calculated according to the density.

### Statistics

All data were presented as mean ± standard error of the mean (SEM). All statistical analyses were performed using GraphPad Prism 9 software (GraphPad Software Inc., San Diego, CA, USA). For neuron morphology and spine and synapse data, values for single cells were averaged for each animal and then per genotype. For the mean apical and basal dendritic length, and all spine and synapse data, a two-tailed Student’s t-test was performed.

## Acknowledgments

We thanks to Alejandro Munizaga and Oscar Ardiles for technical assistance in EM experiments. We also thanks to Enzo Seguel and Leticia Toledo for animal care supervision.

## Author contributions

CF-M, FG-R, MA-P, OS, and AOA designed the research; CF-M, FG-R, MA-P, OS, EM, DL-E, AMG-J and AOA performed experiments, analyzed data and contributed to figures. MF, ADM, and AMG-J, contributed to analytic tools, CF-M, AMG-J, MF, ADM, and AOA wrote the paper.

## Data and materials availability

All data needed to evaluate the conclusions in the paper are present in the paper or the Supplementary Materials.

---

**Fig. S1.**
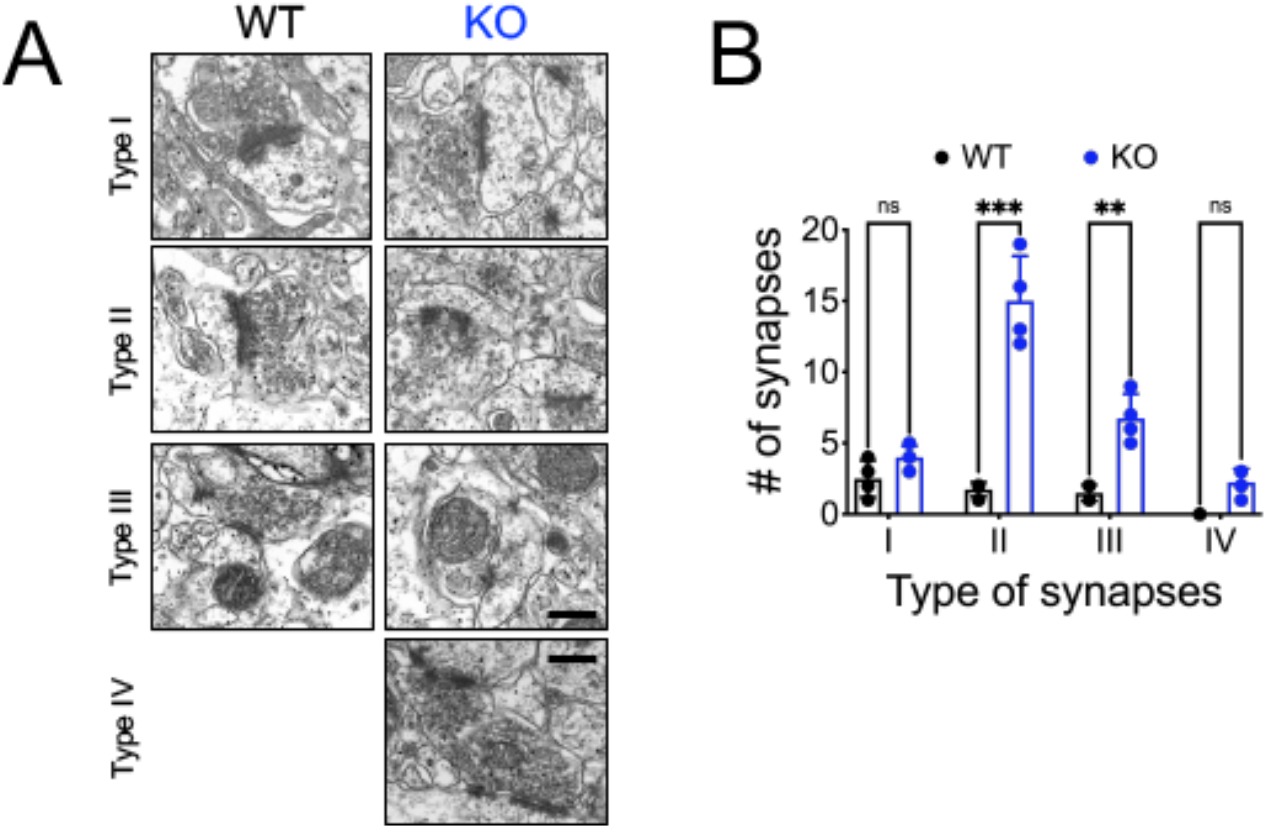
Morphological classification of hippocampal synapses. Representative transmission electron microscopy photographs of asymmetric synapses (**A**) and analysis of the types of synapses (**B**) in CA1 *Stratum radiatum* area of wild type (WT, black) and PANX1-KO (KO, blue) mice. Magnification 43000X, bar: 500 nm.

**Fig. S2.**
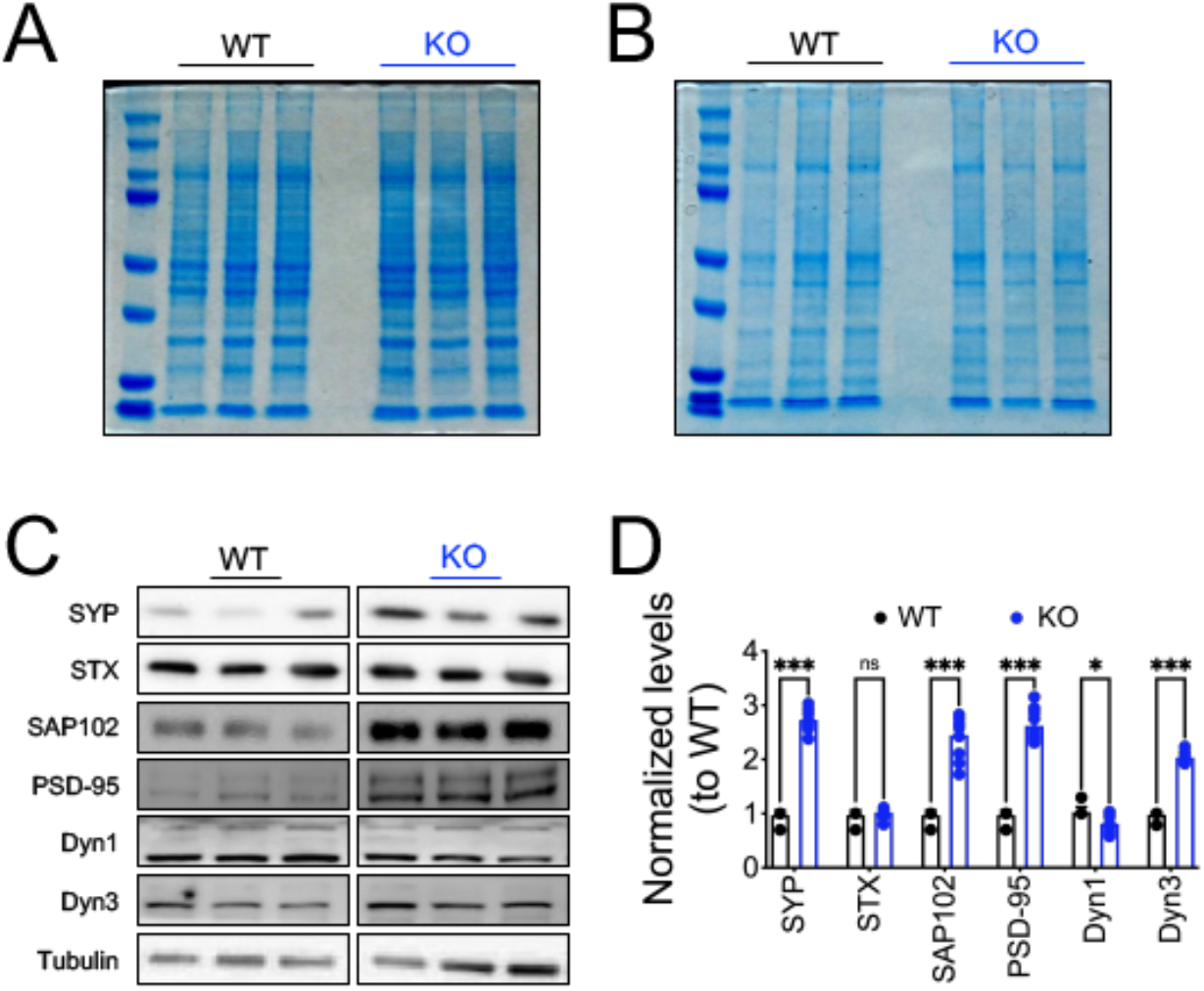
Synaptic proteins levels of hippocampal tissue. Representative images of SDS-PAGE and Coomassie blue-stained gels of hippocampal extracts (**A**) and PSD-enriched fractions (**B**). Representative blots (**C**) and densitometric analysis of hippocampal synaptic proteins.

