## Supplementary figures and images for "Long-term Pannexin 1 ablation promotes structural and functional modifications in hippocampal neurons through the regulation of actin cytoskeleton and Rho GTPases activity"

### Supplemental fig 1

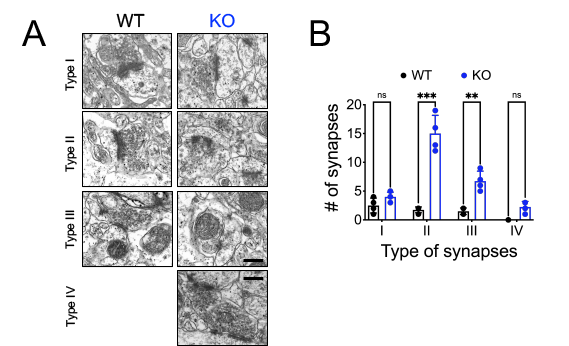

### Supplemental fig 2

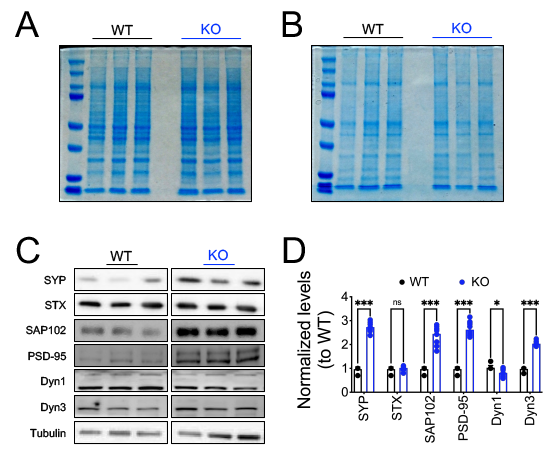
